# Spatial auditory change detection in listeners with hearing loss

**DOI:** 10.64898/2025.12.10.693419

**Authors:** Katarina C. Poole, Simon With, Vincent Martin, Maria Chait, Lorenzo Picinali, Martha Shiell

## Abstract

Everyday listening relies on the auditory system’s ability to automatically monitor the background soundscape and detect new or changing sources. Although change detection is a fundamental aspect of situational awareness, little is known about how hearing impairment affects this ability. This study examined how sensorineural hearing loss influences spatial auditory change detection. Older hearing-impaired listeners (N = 30) completed a spatial change detection task requiring them to identify the appearance of a new sound source within a complex spatialised acoustic scene. Hearing loss was characterised by three factors that were measured with standard clinical tests: audiometric hearing thresholds, sensitivity to small level changes, and sensitivity to spectrotemporal modulation. Simple and mixed-effects linear models were used to test how these factors predicted reaction time, hit rate, and false alarm rate.

Listeners with poorer spectrotemporal sensitivity, higher audiometric hearing thresholds, and older age showed slower and less accurate detection, whereas sensitivity to small changes in level did not predict outcomes. Detection also varied with spatial location, where appearing sources from behind were detected more slowly and less accurately than those from the front or sides. Numerical analysis using head-related transfer functions confirmed that these rear-field effects were unlikely to be explained by overall or frequency-specific acoustic level differences. These findings reveal that hearing loss, age, and spatial factors jointly shape listeners’ ability to monitor dynamic auditory scenes. Additionally, testing spectrotemporal sensitivity offers a promising clinical measure of non-speech auditory processing with relevance for hearing-aid fitting and situational awareness.

## 1. INTRODUCTION

Human listeners are remarkably adept at detecting the emergence of new sound sources, a skill that is essential for navigating and responding effectively in dynamic, unpredictable environments (Aman et al., 2021; Constantino et al., 2012; de Kerangal et al., 2021; Demany et al., 2017; Gregg and Samuel, 2008; Irsik et al., 2016; Pavani and Turatto, 2008; Sohoglu and Chait, 2016). This process, known as auditory change detection, is thought to engage both automatic, pre-attentive mechanisms as well as higher-level attentional systems (Constantino et al., 2012; Eramudugolla et al., 2005; Irsik et al., 2016; Sohoglu and Chait, 2016). It is hypothesised to support situational awareness, with the advantage that it functions even when attention is diverted or visibility is limited, enabling swift responses to novel sounds. Such background monitoring plays an important role in everyday situations such as crossing a street, hearing a familiar voice in a crowd, or catching a train station announcement, and is critical in high-risk environments such as construction sites or battlefields.

Not all listeners benefit equally from the auditory system’s automatic monitoring capabilities. Individuals with hearing loss often report feeling handicapped with behaviours related to situational awareness, such as an impaired ability to judge the direction of approaching footsteps, detect moving vehicles, or localise everyday environmental sounds (Gatehouse and Noble, 2004). Hearing aids can restore access to previously unnoticed environmental sounds, where first-time users often note the return of birdsong or wind (Lelic et al., 2025), yet no quantitative method exists to evaluate how effectively these devices support auditory situational awareness. As auditory change detection is a core mechanism underlying situational awareness, characterising this process in hearing-impaired listeners is critical for understanding how hearing loss disrupts environmental monitoring. A validated measure of auditory change detection in this population would also provide a basis for improving hearing aid design and rehabilitation strategies.

Auditory change detection has traditionally been studied using monophonic pure-tone stimuli, which provide limited insight into how listeners monitor complex acoustic environments (Aman et al., 2021; Constantino et al., 2012). More recent work has shifted toward ecologically-valid paradigms that incorporate spatially distributed sound sources. Poole et al. (2025a) established a benchmark in clinically normal-hearing adults by testing detection of appearing sound sources presented unpredictably across 360° under varying numbers of competing background sources. Spatially separating the sources improved detection, compared to when the sources were co-located, but performance declined as the number of competing sources increased. Importantly, even within this normal-hearing group, elevated mid-to-high frequency thresholds (2–16 kHz) predicted poorer performance, an effect observed only in spatialised conditions, suggesting that subtle auditory deficits can limit spatial monitoring. EEG recordings confirmed robust neural markers of change detection despite passive listening and the complexity of the stimulus set. Interestingly, in a separate behavioural task, button-press responses were slower for sounds appearing behind or above the listener, than those from frontal or lateral positions. The authors attributed these asymmetries not to peripheral acoustic filtering but to later-stage attentional or decision-making processes, since ERP amplitudes did not differ by source location, although this interpretation remained untested.

Building on this benchmark, the present study examined how hearing loss alters spatial auditory change detection, using the paradigm employed by Poole et al. Given that Poole et al. showed even subtle high-frequency deficits constrain performance in normal-hearing adults, it is reasonable to expect that the elevated hearing thresholds characteristic of clinically diagnosed sensorineural hearing loss would further impair individuals’ ability to detect appearing sound sources in a scene. Evidence from related contexts support this expectation, for example when hearing level detection thresholds are artificially elevated through the use of protective equipment in noisy work environments, adults with otherwise normal hearing show delayed detection of threatening sounds (Clasing and Casali, 2014), and report impairments to situational awareness (Abel, 2008; Reddy et al., 2014).

Beyond reduced audibility, individuals with hearing loss often experience additional distortion effects that degrade the clarity of sounds (Plomp, 1978) and hinder auditory object detection in noise (Boncz et al., 2024). Notably, suprathreshold hearing-in-noise deficits can persist even when cochlear amplification is accounted for (Johannesen et al., 2016; Peters et al., 1998). For listeners with sensorineural hearing loss, this hearing-in-noise deficit is often attributed to a loss of frequency selectivity and inability to use temporal fine structure information to separate competing sounds into separate streams (Bernstein et al., 2013; Lorenzi et al., 2006). Such deficits would be expected to impair auditory change detection in complex environments, even if hearing levels are compensated for.

Paradoxically, it is also possible that sensorineural hearing loss could support improvements to auditory change detection through the loss of the cochlea’s compression function. In healthy cochleas, the outer hair cells respond to sound level non-linearly with compression at medium levels (e.g. Robles et al., 1986). A loss of this compression functionality in people with cochlear damage can lead to an increased sensitivity to loud sounds, and improved detection of changes in sound level (see for reviews: Buus et al., 1982; Moore, 1996). Potentially, in a change detection task, this loudness recruitment could offset some of the perceptual challenges posed by a loss of audibility and impaired hearing-in-noise ability.

In the current study, we tested for a relationship between individual’s performance in spatial change detection and three clinical measures of their hearing loss. First, we assessed listeners’ hearing-in-noise ability using the Audible Contrast Threshold (ACT) test, a clinical tool that measures spectrotemporal modulation detection thresholds (Zaar et al., 2024b), and is predictive of a listener’s ability to hear speech-in-noise (Zaar et al., 2023). Second, we assessed listeners’ sensitivity to small level changes using the Short-Increment Sensitivity Index (SISI) test (Jerger et al., 1959). Clinically, this test is meant to distinguish cochlear from retrocochlear etiologies (Ruggero and Rich, 1991; Shi et al., 2022), where high scores are associated with the former, due to a loss of the compression functionality of the outer hair cells. Given our participant recruitment criteria, we expected that all participants would meet the criteria for cochlear damage, and we rather used their SISI scores to capture individual differences in sensitivity. As the ACT and SISI tests were the focus of our investigation, it was necessary to ensure that participants were able to hear the stimuli despite their hearing loss. As such, we adjusted the sound levels for each participant based on their audiogram, following the NAL-RP linear gain rule (Dillon, 2012). We nonetheless included hearing threshold levels (HTL) as a third clinical measure, to account for potential remaining effects of decreased audibility.

With this research, we aimed at addressing two main research questions. First, how does spatial auditory change detection vary with different features of hearing loss? We predicted that listeners with poorer hearing, characterised by elevated HTLs and ACT scores, would show slower reaction times and reduced detection accuracy, whereas higher SISI scores might offer an advantage by enhancing sensitivity to level increases from the appearing source. Second, does spatial auditory change detection vary with source location in hearing-impaired listeners? Based on prior findings in normal-hearing adults, we expected performance to be worse for sounds emerging from behind the listener. To further probe the basis of this effect, we also carried out numerical analyses to determine whether the observed differences could be explained purely by acoustic factors. By integrating clinical measures with a controlled spatial paradigm, this study aimed to clarify how hearing loss shapes the spatial dynamics of auditory attention in complex environments.

## 2. MATERIAL AND METHODS

### 2.1 Participants

Participants were recruited from the Eriksholm Research Centre test participant pool, where they received audiological services and hearing aids in exchange for their membership. They were financially compensated for travel to the test site at Eriksholm Research Centre. The research abided by established international research codes as per the declaration of Helsinki. A total of 32 adults with hearing loss were tested; however, one participant was excluded due to difficulty in understanding the change detection task and another because an ACT score could not be obtained. The final sample comprised of 30 adults (16 males and 14 females, mean age 69 years, standard deviation 11 years; see Figure 3A for distribution). The overall 4-frequency pure-tone average (PTA) across all participants was 40.7 dB HL across both ears, with audiograms generally falling within the N2-N5 moderate sloping range (Bisgaard et al., 2010). Hearing threshold levels were symmetrical within a range of 15 dB HL in at least three out of the frequencies 0.5, 1, 2 and 4 kHz. The group average and individual hearing thresholds and asymmetry are illustrated in Figure 1; extended high-frequency hearing thresholds are presented in Table 4 of the Supplementary material. All participants had adult-onset hearing loss with presumed age-related origin, except one who additionally had a childhood-onset hearing loss. Table 4 All participants were native Danish speakers with no prior exposure to the testing materials and gave informed written consent. While they were experienced hearing aid users, they completed the experiment without their hearing aids.

**Figure 1:**
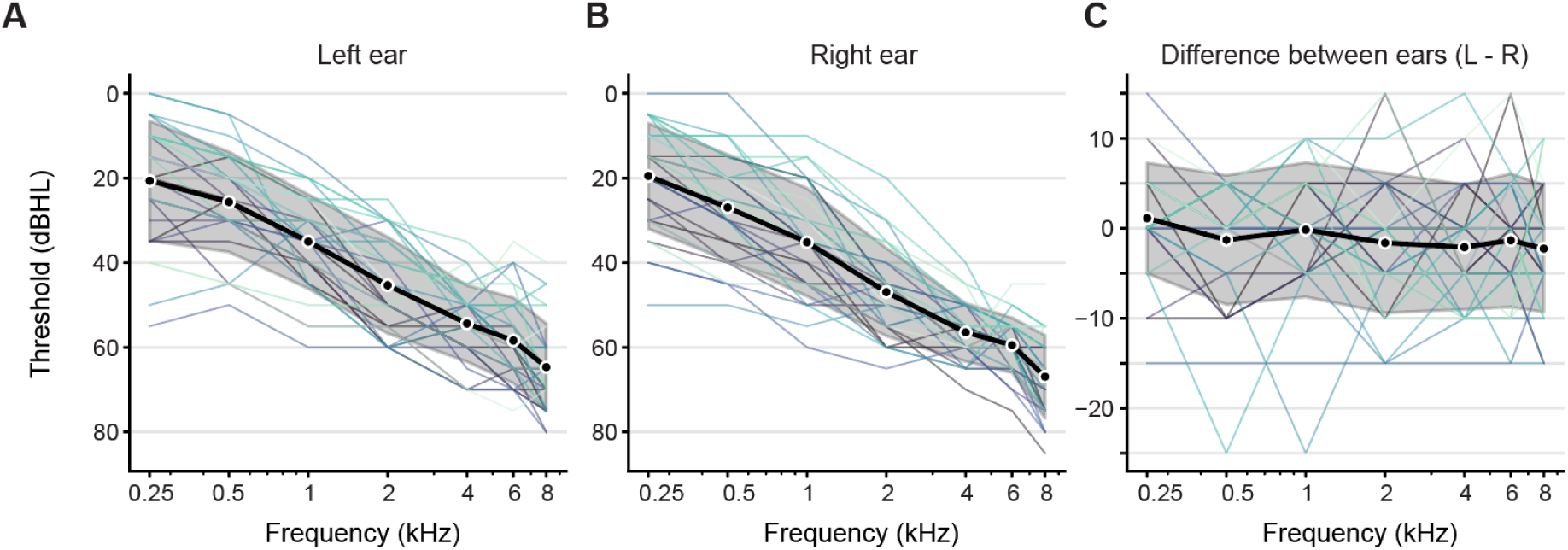
Participant audiograms. Audiograms are shown between 0.25 kHz and 8 kHz for each participant (coloured lines) as well as the group average (black line) and standard deviation (shaded region) for the left ear (A) and right ear (B). C) Difference in hearing threshold between the left and right that show any asymmetry in hearing loss for each participant and across the participant pool.

### 2.2 Procedure

#### 2.2.1 Clinical measures

All clinical tests were administered by an experienced audiologist with an Affinity Compact audiometer (Interacoustics model AC440) in a sound-treated booth. Stimuli were presented via Radio Ear headphones (model DD65), and participant responses were recorded with a thumb-trigger. The participants were first examined with standard otoscopy and pure-tone audiometry at 0.25, 0.5, 1, 2, 4, 6, 8, 10, 12, and 16 kHz. The extended high-frequency thresholds (from 8-16 kHz) were collected to describe our test participants, since elevated HTL in high-frequencies is associated with impaired spatialised change detection performance in people with clinically-normal hearing thresholds (Poole et al., 2025a). Considering the numerous instances where thresholds were at ceiling level (see Table 4 in Supplementary), we chose not to include the extended high-frequency values in our analysis. As such, for each participant, HTL were calculated as the average HTL from 0.25 to 6 kHz in both ears.

After audiometry, participants completed the ACT and SISI tests, following the standard implementation in the Affinity Compact audiometer. The ACT task uses a staircase procedure, controlled in real-time by the audiologist, to measure a listener’s ability to detect a spectrotemporal modulation of pink noise presented to both ears simultaneously. Scores are provided on a normalized contrast level (nCL) scale ranging from –4 to 16, where lower values indicate better hearing-in-noise, and a score of 0 is the expected median level for young, normal-hearing listeners (Zaar et al., 2024b). In the SISI task listeners were asked to detect a 1 dB change in the level of a continuous pure-tone presented at 20 dB (HL) above HTL. Each ear was tested separately with twenty level changes played automatically at unpredictable timepoints, for each frequency tested. All participants were tested at 4, 2, and 1 kHz. If participants had responses at 1 kHz, they were additionally tested at 0.5 kHz, otherwise they were assigned a score of 0 for this frequency. We calculated each participants’ percent correct detections across the four frequencies in both ears.

#### 2.2.2 Apparatus

For both the reaction time (RT) baseline measurement and the scene detection task, the participant sat in the chair in the middle of a 24-loudspeaker ring in a semi-anechoic room. The loudspeakers (Genelec 8030A and 8030C) were fixed from the ceiling at the approximate height of the test participant’s ears and distributed at 15-degree intervals (circle diameter: 2.4 m). A 29 x 16.5 cm screen was mounted on an adjustable stand next to the chair so that it could be positioned in front of the participant’s face, with an approximate distance of 65 cm. Max/MSP software (Cycling74, version 8.1) managed stimulus delivery via a Dante controller connected via Ferrorfish and Yamaha sound cards, triggered by custom Python scripts (Python Software Foundation, version 3.8) that also provided text prompts and visual stimuli to the screen in front of the participant. Participant responses were recorded via a Bluetooth-connected number pad. Participants were instructed to hold the number pad in their non-dominant hand and make their responses by pressing “0” with the thumb of their dominant hand. They were presented with a fixation cross throughout the experiment and were asked to keep their eyes open and to look at the cross.

#### 2.2.3 Reaction time baseline measurement

To account for individual differences in auditory-motor reactivity, we measured baseline reaction times in response to a sound in silence. For this measurement, the participants were told that a sound would be played from the loudspeaker directly in front of them, and that they should press the response button as soon as possible when they heard it. In each trial, a sound was randomly selected from the stimuli that were used in the change detection task (described below). These were presented with a time interval that varied randomly between 1 to 5 seconds. After the participants’ response, their RT was presented on the screen as feedback for 2 seconds, after which the next trial began automatically. Participants completed 48 trials in total.

#### 2.2.4 Change detection task

We tested the ability of the participants to detect an appearing sound source and how the location of the appearing sound source influenced their performance. The stimuli used were identical to those used in Poole et al. (2025) and consisted of nine-second-long broadband sounds with natural temporal envelopes and complex spectral content, designed to retain real-world acoustic complexity whilst removing semantic information(see Figure 2A-B). Envelopes were extracted from natural sounds (e.g., vocalisations, speech babble) and applied to carriers with varied harmonic and noise characteristics. The initial sound pressure level for each sound was set to 62 dBA, as measured from the approximate position of the listener’s head in the centre of the loudspeaker array. Prior to the start of each session, stimuli levels were adjusted for each individual based on their audiogram, using NAL-RP linear gain rule (Dillon, 2012). This adjustment was intended to compensate for impaired audibility of the stimuli. To reduce the risk that the sounds became uncomfortably loud for the participants, we limited this adjustment to frequencies between 0.25 and 6 kHz. Participants were also given an opportunity to lower the overall level of the stimuli during the training.

**Figure 2:**
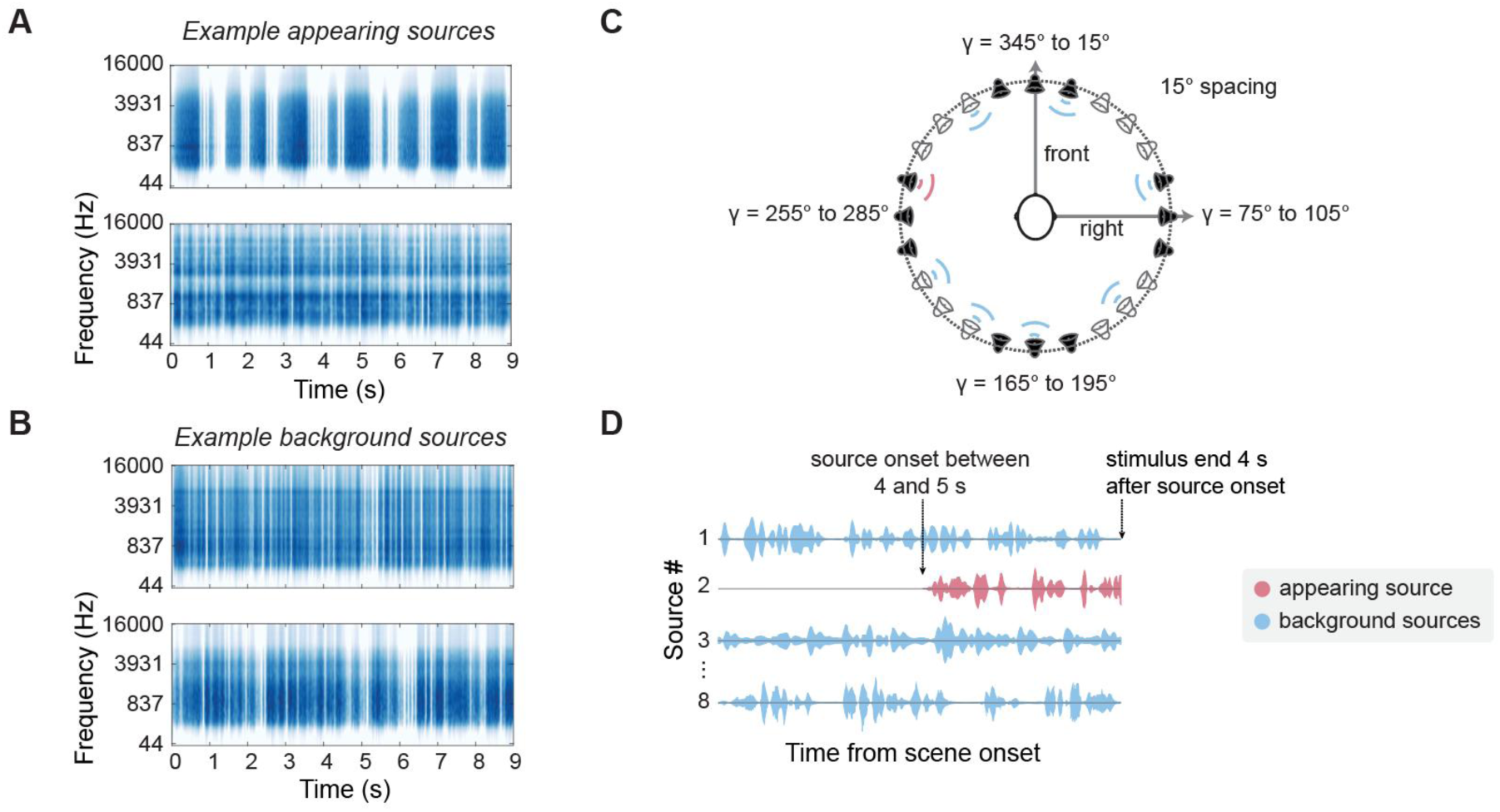
Acoustic stimuli and scene structure and apparatus. ERB-weighted spectra of two examples of appearing sources (A) and two examples of background sources (B). C) Horizontal loudspeaker array layout used. The participant was situated at the centre of the sphere. The array consisted of 24 loudspeakers separated in 15-degree intervals. Appearing sound sources could come out of one of three speakers at the front, right, back or left (filled in black in the figure). An example source location layout for a trial is also displayed where the pink waves in front of the speaker denote the appearing source and the blue waves the background sources. D) Example schematic of the acoustic scenes used with concurrent sources and an appearing sound source 4 to 5 seconds from the onset of the acoustic scene. All sources overlapped in frequency space and the whole stimulus ended 4 seconds after appearing source onset. In catch trials (“no change”) there was no appearing sound source and the whole acoustic scene lasted between 8 and 9 seconds.

The scene stimuli were presented over the 24-loudspeaker array (see Figure 2C) in a semi-anechoic room. The trial began with a scene of seven different sounds (“background sources”) playing simultaneously from pseudo-randomly selected locations around the listener, such that at least one sound, and no more than two, appeared in each quadrant. The background sources were randomly selected from the 11 possible stimuli. In half the trials, a new sound source would appear between 4 and 5 seconds from scene onset, making eight concurrent sources (see Figure 2D). The participants were told to react to the onset of the appearing source by pressing the response button on the number pad as quickly as possible, but only if they heard the appearing sound source. We measured the participants’ reaction time and accuracy in detecting the appearing source whilst changing the location of the sound source. The location of the appearing sound source was restricted pseudo-randomly to one of 12 loudspeakers, either in the front (345 to 15), left (255 to 285), right (75 to 105) or back speakers (165 to 195), such that each of these 12 loudspeakers was used for an appearing sound source twice in each experiment block. This totalled to 6 trials for each of the four location conditions per experiment block. Trials were presented in three blocks of 48 trials each (approximately seven minutes). Eighteen repeats were presented per location condition with 72 “no change” trials. Within a block, an inter-trial-interval of 2 s was used.

After the participant’s response, the stimuli were stopped. If no response was made, the stimuli continued for 8 seconds. At the end of each trial, participants were informed of their accuracy and their reaction time, if applicable, in the previous trial (i.e. no RT information was given if the participant made an error or did not make a response). The feedback was displayed for 2 seconds, and then the next trial began automatically. Participants completed at least one block of 48 practice trials, during which they were given the opportunity to ask for clarification if needed. This was followed by three blocks of 48 trials, each of which took approximately 7 minutes to complete. Participants had the opportunity to take a break in between blocks, and the experiment took between 1.5 to 2.25 hours in total to complete.

### 2.3 Behavioural analysis

For each participant, three independent variables (IVs) were derived from the clinical tests: ACT scores, average hearing threshold levels, and SISI scores. To account for shared variance between the clinical predictors and avoid multicollinearity in the model, we calculated residual ACT and residual SISI scores. This was done by regressing each score (ACT and SISI) against HTL and extracting the residuals from those regressions. The resulting residual scores represent the variance in ACT and SISI performance that is independent of hearing threshold levels. This allowed us to isolate the unique contribution of listeners’ ability to analyse fine sound features (as captured by ACT and SISI scores) from the influence of basic hearing sensitivity (HTL), thereby improving the interpretability of our results.

To address our first research question, identifying which clinical scores predicted change detection performance, we took a model building approach. For this, we constructed three different linear models that each predicted one of three measures of scene detection performance (i.e. the dependent variables): average RT to correctly detected appearing sources, hit rate, and false alarm rate (see Figure 3A-E). The average RT for correctly detected appearing sources was calculated as the log inverse of the RTs in the scene detection task. Note that with this transformation, faster (smaller) RTs are expressed as larger values, and RTs greater than 1 second are expressed as negative values. Hit rate and false alarm rate were analysed independently to align the results with our second research question on how source location impacts change performance.

**Figure 3:**
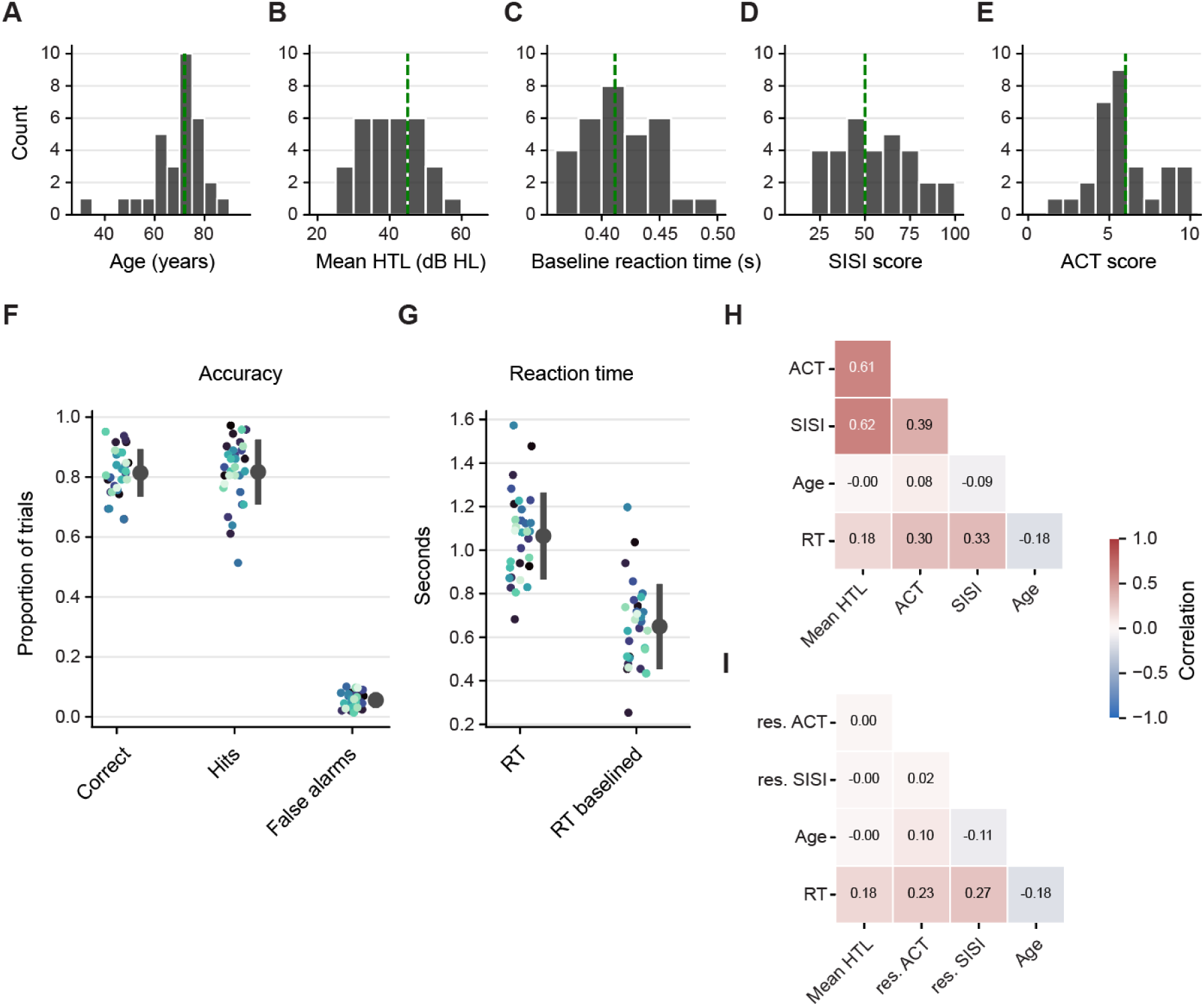
Clinical and behavioural measures of participants. A–E) Distributions of participant characteristics and median in dashed green, including (A) age, (B) hearing threshold level (HTL, averaged across ears and frequencies), (C) baseline reaction time, (D) Short Increment Sensitivity Index (SISI) scores, and (E) Audible Contrast Threshold (ACT) scores. F) Behavioural accuracy for each participant (coloured dot) and mean and standard deviation across participants for the proportion of correct trials, hits and false alarms. G) Reaction time for correct detections and baselined reaction time. H) Correlation matrix showing relationships among clinical measures and covariates, with warmer colours indicating stronger positive correlations. I) Correlation matrix of the residual ACT and SISI scores after regressing out shared variance with HTL, used as predictor variables in subsequent analyses.

The independent variables tested in each model were HTL, SISI score residuals, and ACT score residuals. These were transformed into z-scores to facilitate interpretation of the calculated coefficients relative to each other. For predicting RT on the scene detection task, we included the log inverse baseline RT as a covariate. Likewise false alarm rate was included as a covariate for predicting hit rate, and hit rate was used as a covariate for predicting false alarm rate. Age was always included as a covariate. Our model-building approach used a stepwise elimination procedure based on p-values, with a cutoff for exclusion set at p > 0.1. At each step, the procedure began with an initial model. The IV with the highest p-value exceeding the cutoff was removed to create a new model. Each previously excluded IV was then reintroduced individually to ensure it still met the exclusion criterion; if not, it was reinstated. This updated model became the new initial model for the next step. The process repeated until all remaining IVs had p-values below the exclusion threshold.

To address our second research question, we compared two linear mixed models to determine whether spatial location improved predictions of detection performance using a likelihood ratio test. The “Location model” included spatial location as an IV along with relevant covariates, while the “Covariates-only model” included only the covariates identified in the first analysis. In both, participant identity was included as a random effect. Two dependent variables were tested: (1) average reaction time (log-inverse transformed) and (2) hit rate, both calculated separately for each location. False alarm rates were excluded, as these trials were not associated with a specific location.

### 2.4 Numerical acoustic analysis

To verify playback conditions and assess whether acoustic factors such as head shadowing or filtering could help explain the behavioural results, we conducted a numerical simulation in which the free-field scenes were rendered binaurally. To achieve this, trials were defined as that of the change detection experiment and rendered binaurally using individual head-related transfer functions (HRTFs). We randomly selected 100 HRTFs from the SONICOM HRTF dataset (Engel et al., 2023; Poole et al., 2025b) to ensure as wide a range of morphologies were used. Python modules py3dti, spatialaudiometrics, and scipy were employed to render the binaural stereo audio files and perform acoustic analysis (Pauwels, 2023; Poole et al., 2025b). To assess whether overall level cues might have influenced performance, we examined changes in root-mean-square (RMS) levels before (−600 to 0 ms) and after (0 to 600 ms) target onset on trials containing an appearing source. Such cues could potentially allow perceptual differentiation between source locations or reduce performance in conditions where the appearing source was quieter. RMS was computed separately for the left and right ears and averaged across ears, for each condition and individual HRTF. This was also calculated as a function of frequency by passing the left and right signal through a gammatone filter bank spanning 20 Hz to 20 kHz with 32 Equivalent Rectangular Bandwidth (ERB) centre frequencies logarithmically spaced. The change in RMS before and after target onset was then calculated for each frequency band for each individual.

## 3. RESULTS

### 3.1 Clinical measures predict change detection performance

Our first research aim was to examine how spatial auditory change detection varies with hearing loss, focusing on whether audiometric thresholds and clinical tests of auditory processing can explain individual differences in detection performance. To address this, we characterised participants (N = 30) using a range of clinical and demographic measures: audiometric hearing threshold levels (HTL), Short Increment Sensitivity Index (SISI) scores, Audible Contrast Threshold (ACT) scores, and age. Distributions of these measures are shown as histograms in Figure 3A-E). The average participant age was 68.9 years (SD = 11.27), and the mean HTL was 40.2 dB HL (SD = 8.4). SISI scores ranged from 23.75 to 97.5 (mean = 54.41), and ACT scores ranged from 1.9 to 9.5 (mean = 5.77), reflecting considerable variation spectrotemporal sensitivity. Behaviourally, participants performed the change detection task with good accuracy (see Figure 3F-G). The mean proportion of correct trials was 0.814 (SD = 0.0792), with a low false alarm rate of 0.057 (SD = 0.0256) and a mean hit rate of 0.817 (SD = 0.108). Reaction times for correct detections averaged 1.065 s (SD = 0.199), and after baseline correction 0.649 s (SD = 0.195), establishing a robust behavioural baseline against which the influence of hearing thresholds and auditory processing measures could be assessed.

Correlations among the independent variables and covariates are shown in Figure 3H, where moderate correlations were observed between SISI scores, and ACT scores against HTL (0.62 and 0.61 respectively). The association between ACT and HTL is consistent with previous findings that more severe hearing loss is linked to reduced sensitivity to complex sound features (Zaar et al., 2024a). Similarly, the relationship between SISI scores and HTL reflects the known pattern in sensorineural hearing loss, where damage to outer hair cells supports enhanced SISI performance which we observed in our participant group. To isolate the independent effects of ACT and SISI, we regressed each on HTL and used the residuals as predictors (see Figure 3I). These residual scores reflect performance on the ACT and SISI after accounting for hearing threshold differences, allowing assessment of their unique influence on spatialised change detection.

To understand whether individual differences in hearing impairment can impact spatial auditory change detection, we examined whether individual differences in HTL, ACT, SISI, age or baseline reaction time could account for variation in three dependent measures from the change detection task: reaction time (RT), hit rate, and false alarm rate. To do this we used the model-building approach described in section 4.3 (Analysis). Table 1 reports the coefficients and associated p-values from the final models for each outcome. ACT residuals, HTL, and age were all significant predictors of both reaction time and hit rate. Participants with better ACT scores (lower scores) not only responded more quickly (higher log-inverse RTs) but also detected more newly-appearing sources, suggesting that sensitivity to fine-grained acoustic structure supports more efficient change detection. Descriptively, the estimated coefficients for HTL and ACT residuals were of similar magnitude, indicating that the effect of these factors on performance was similarly weighted. We further observed that older participants tended to respond more slowly and with lower accuracy. Age was also the only significant predictor for the false alarm rate, with older participants displaying a higher false alarm rate. The baseline reaction time significantly predicted reaction time to the change detection task where quicker baseline reaction times led to quicker change detection reaction times, indicating that general response speed influenced task performance. However, across response measures SISI scores did not emerge as a significant predictor.

**Table 1:** Final model coefficients for each behavioural outcome measure: false alarm rate, hit rate and reaction time (log-inverse). Only predictors that were retained in the final models following the model-building procedure are shown. The standard error, T and p values are also displayed for each significant predictor. False Alarm Rate model: Residual standard error = 0.0939 on 29 degrees of freedom, Multiple R^2^ = 0.190, Adjusted R^2^ = 0.162, F(1,29) = 6.804, p-value: 0.0142. Hit Rate model: Residual standard error = 0.0787 on 26 degrees of freedom; Multiple R^2^ = 0.532, Adjusted R^2^ = 0.479, F(3,26) = 9.871, p-value < 0.001. Reaction time model: Residual standard error = 0.114 on 25 degrees of freedom; Multiple R^2^ = 0.560, Adjusted R^2^ = 0.489, F(4,25) = 7.946, p-value < 0.001.

| Fixed effects | Estimate | Standard error | T | p value |
| --- | --- | --- | --- | --- |
| *False Alarm Rate* |  |  |  |  |
| **Intercept** | -0.051 | 0.106 | -0.477 | 0.637 |
| **Age** | 0.004 | 0.002 | 2.608 | **0.014** |
| *Hit Rate* |  |  |  |  |
| **Intercept** | 1.071 | 0.089 | 11.971 | **< 0.001** |
| **HTL** | -0.050 | 0.014 | -3.508 | **0.002** |
| **ACT (residual)** | -0.040 | 0.015 | -2.698 | **0.012** |
| **Age** | -0.004 | 0.001 | -2.900 | **0.007** |
| *Reaction time (log inverse)* |  |  |  |  |
| **Intercept** | -0.203 | 0.346 | -0.586 | 0.563 |
| **HTL** | -0.044 | 0.021 | -2.098 | **0.046** |
| **ACT (residual)** | -0.078 | 0.022 | -3.538 | **0.002** |
| **Age** | -0.006 | 0.002 | -2.980 | **0.006** |
| **Baseline RT (log inverse)** | 0.714 | 0.327 | 2.183 | **0.039** |

### 3.2 Location of the appearing source impacts performance

Our second research aim was to determine whether spatial auditory change detection varies with source location in hearing-impaired listeners, particularly for sounds emerging from behind the listener. To test this, we examined whether reaction times and hit rates were influenced by the spatial location of the appearing source, beyond the effects of individual differences in hearing and auditory processing. Using a model comparison approach (see section 4.3, Analysis), we found that including the spatial location of the newly appearing source significantly improved prediction of both reaction time (RT) (ΔAIC = 33.9, χ²(3) = 39.9, p < 0.0001) and hit rate (ΔAIC = 6.3, χ²(3) = 12.3, p < 0.01), beyond models containing only participant covariates: age, ACT residuals, HTL, and baseline RT (for RT models). For RT, the combined model with location and covariates explained 62.9% of the variance, compared to 43.5% explained by covariates alone. For hit rate, location and covariates together explained 52.3% of variance, versus 46.5% explained by covariates alone.

Model coefficients and associated p-values are presented in Table 2 and Table 3 for reaction time and hit rate respectively. Estimate marginal means are shown in Figure 4A-B, with the back location set as the reference level in the model for location. Consistent with our hypothesis, reaction times were slower for newly-appearing sounds originating from behind participants relative to sounds from the front, left, or right. No significant reaction time differences were observed among the front, left, and right locations. Similarly, hit rates were lower for sounds originating from behind compared to those from the front and left. However, no significant difference was found between back and right locations, potentially reflecting reduced detection performance for sounds from the right, which showed a lower hit rate compared to the front. No significant difference was observed between left and right locations.

**Figure 4:**
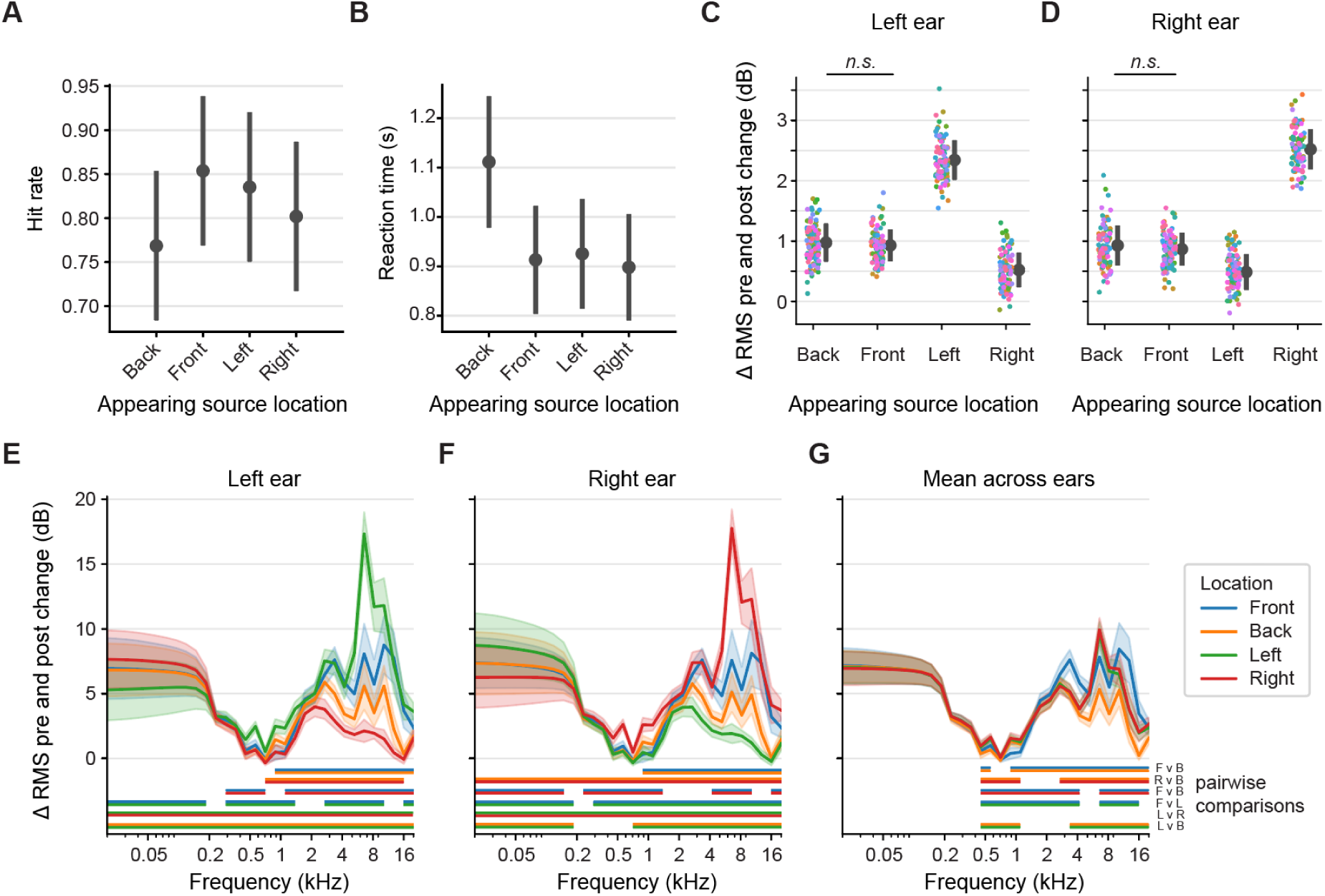
Effect of appearing source location on performance and simulated acoustic cues. (A–B) Estimated marginal means of hit rate (A) and reaction time (RT) (B) as a function of appearing source location, derived from the mixed-effects models. Error bars show the 95% confidence intervals on the estimated marginal means (C–D) Simulated change in root mean square (RMS) level at each ear (ΔRMS) pre-versus post-source onset, based on head-related transfer functions (HRTFs) from 100 individuals. (E–F) Frequency-dependent ΔRMS across ERB-weighted frequency bands for the left (E) and right (F) ears. (G) Mean binaural ΔRMS across ears. (E–G) Lines are coloured by source location; shaded areas show ±1 SD. Significant pairwise differences (cluster-based permutation tests) are indicated below, with colour pairs denoting the compared source locations.

**Table 2:**
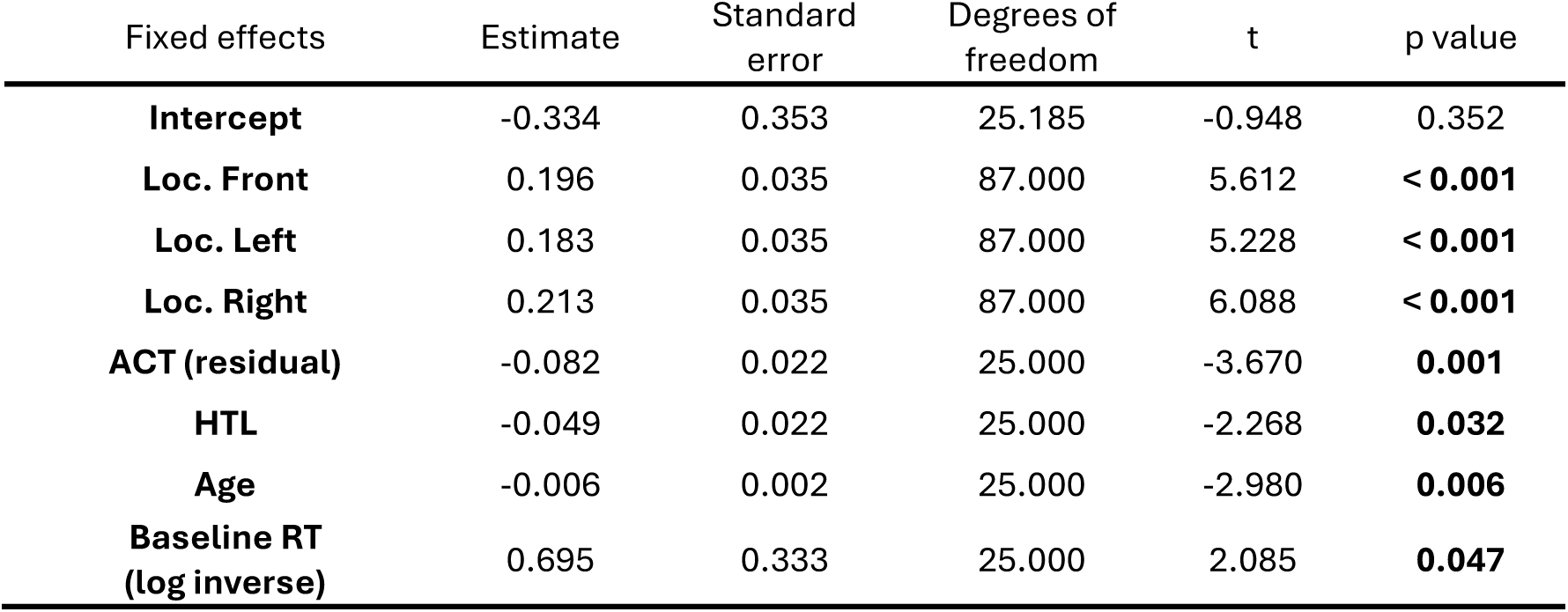
Fixed-effect coefficients from the final linear model predicting reaction time (log-inverse) as a function of source location and participant covariates. The model includes spatial location (back = reference level, “Intercept”), ACT residuals, hearing threshold levels (HTL), age, and baseline reaction time. Estimates, standard errors, degrees of freedom, t-values, and p-values are reported for each predictor. Marginal R^2^ = 20.451 and Conditional R = 0.629.

**Table 3:** Fixed-effect coefficients from the linear mixed-effects model predicting hit rate as a function of source location and participant covariates. The model includes spatial location (back = reference level, “Intercept”), ACT residuals, hearing threshold levels (HTL), and age. Estimates, standard errors, degrees of freedom, t-values, and p-values are reported for each predictor. Marginal R^2^ = 20.346 and Conditional R = 0.523.

| Fixed effects | Estimate | Standard error | Degrees of freedom | t | p value |
| --- | --- | --- | --- | --- | --- |
| **Intercept** | 0.768 | 0.021 | 88.175 | 35.982 | **< 0.001** |
| **Loc. Front** | 0.085 | 0.026 | 87.000 | 3.306 | **0.001** |
| **Loc. Left** | 0.067 | 0.026 | 87.000 | 2.587 | **0.011** |
| **Loc. Right** | 0.033 | 0.026 | 87.000 | 1.293 | 0.199 |
| **ACT (residual)** | -0.040 | 0.015 | 26.000 | -2.698 | **0.012** |
| **HTL** | -0.050 | 0.014 | 26.000 | -3.508 | **0.002** |
| **Age** | -0.042 | 0.014 | 26.000 | -2.900 | **0.007** |

To determine whether the observed delays in reaction time and reduced performance for sounds appearing behind the listener could be explained by acoustic shadowing effects caused by the head and ears, we conducted simulations using acoustically measured head-related transfer functions (HRTFs) from the SONICOM database (Engel et al., 2023; Poole et al., 2025b). We used HRTFs of 100 randomly selected participants to allow us to model the acoustic input at each ear for all loudspeaker positions and sound source locations. The change in RMS level from pre- and post-appearing source onset was approximately 1dB for the back and front locations for each ear and varied between 0.5 and 2.5 dB for the left and right locations for each ear (see Figure 4C-D). Analysis revealed a significant main effect of source location on the root mean square (RMS) sound level at the ear. Post-hoc pairwise comparisons showed no significant difference in RMS level between the back and front locations, while significant differences were found for the left and right locations, likely reflecting higher sound levels at the ipsilateral ear. This indicates that overall differences in RMS level caused by acoustic shadowing do not explain the longer reaction times or lower detection rates for sounds originating from behind compared to those from the front.

To test whether frequency-dependent acoustic shadowing might explain the behavioural effects, we examined changes in RMS level across ERB frequency bands for each ear and for the binaural average (see Figure 4E–G). Cluster-based permutation statistics revealed significant front–back differences across broad high-frequency ranges (883–20,000 Hz) in both ears, with average differences of 1.585 dB (left ear, p = 0.0059) and 1.606 dB (right ear, p = 0.0059). For the binaural average, a similar high-frequency effect was observed (mean difference = 1.595 dB, p = 0.0059), as well as a narrower low-frequency effect from 452–565 Hz (mean difference = 0.271 dB, p = 0.0234; see Supplementary Table). Within these significant ranges, the largest differences occurred around 12.8 kHz, reaching 5.540 dB (left ear), 5.394 dB (right ear), and 5.467 dB (both ears). These findings quantify the systematic level differences between front and back appearing source locations are present across specific frequency bands.

## 4. DISCUSSION

This study investigated how hearing impairment shapes listeners’ ability to detect changes in complex spatial auditory scenes. Using a multi-source change detection paradigm adapted from Poole et al. (2025a), we addressed two main questions: how performance varies with features of sensorineural hearing loss, and whether detection differs depending on the spatial location of the newly appearing sound. Our findings provide clear evidence on both fronts. First, poorer spectrotemporal sensitivity and higher hearing threshold levels, as indexed by the ACT test and pure-tone audiometry, respectively, were associated with slower reaction times and lower detection accuracy. This was true even after the shared variation between these two measures was accounted for, highlighting the ACT test as a particularly sensitive measure of functional auditory processing, capturing individual differences that extend beyond standard audiometric thresholds. Age also exerted a consistent influence, with older listeners responding more slowly, missing more changes, and making more false alarms. Second, detection performance was reduced for sounds emerging from behind the listener where reaction times were slower and hit rates lower compared to sounds from the front or side. Importantly, HRTF simulations indicate that the observed rear-field deficits are unlikely to be explained by overall or frequency-specific RMS level differences due to acoustic shadowing, suggesting instead that attentional factors limit detection of changes in the rear field. To our knowledge, this is the first study to characterise spatialised auditory change detection in hearing-impaired listeners, providing new insight into how hearing loss affects perception of dynamic acoustic scenes.

### 4.1 Comparison with normal-hearing listeners

A useful point of comparison comes from previous work that employed a similar spatial change detection paradigm in normal-hearing listeners, but with sources distributed in both azimuth and elevation (Poole et al., 2025a). While such data cannot provide a direct benchmark for the present study, given the differences in listener population and task design, they nonetheless offer context for evaluating performance in hearing-impaired listeners, where no prior data is available. In the normal-hearing (NH) study, detection accuracy for scenes containing eight sources was approximately 80% correct, closely matching performance observed here. Variability was also comparable, with a standard deviation of 7.92% in the present study and approximately 11% in the NH group.

Reaction times showed some differences where in the NH study, frontal source detections averaged around 800 ms, and in the current data even after baseline correction, front-source reaction times were closer to 900 ms. However, direct comparison is complicated by methodological differences such as the absence of RT baseline correction in the NH study and inclusion of elevation cues in the NH paradigm which may have enhanced perceptual segregation. These factors make it difficult to draw firm conclusions about reaction time differences between listener groups from the available data. Overall, detection accuracy appears relatively comparable across normal-hearing and hearing-impaired groups when frequency thresholds are compensated for, though further work with matched methodologies would be needed to establish whether reaction times differ systematically between populations.

### 4.2 Clinical measures can be used to assess non-target hearing

Audiometric hearing threshold levels (HTLs) are the standard clinical measure of hearing loss, providing the foundation for both diagnosis and intervention planning. In the present study, HTLs emerged as a significant predictor of both reaction time and hit rate in the change detection task. However, while HTL explained substantial variance, it did not fully account for performance differences across listeners. Additional factors, such as sensitivity to spectrotemporal modulation (ACT scores), contributed to performance. In our study, poorer ACT scores were associated with lower hit rates and slower reaction times in the change detection task, indicating that listeners’ ability to track spectrotemporal modulations supports successful and rapid detection of changes in spatial auditory scenes.

The ACT test is particularly relevant in this context, as it assesses hearing-in-noise ability using non-speech stimuli, in contrast to the common approach of measuring hearing-in-noise performance is through tasks that use speech as either the target or the masker (e.g. Billings et al., 2023; Soli and Wong, 2008). However, speech-based tests may not generalise well to non-speech listening contexts such as the present spatial change detection paradigm. Speech and non-speech sounds engage partially distinct auditory processes (Surprenant and Watson, 2001), and previous work has shown little correlation between speech-in-noise and non-speech-in-noise tests (Vermiglio et al., 2019). This suggests that performance on one type of task does not necessarily predict ability on the other, and that different listeners may experience distinct challenges depending on the acoustic context. The ACT test may provide a valuable bridge between these domains as it offers a language-independent measure of spectrotemporal sensitivity that captures performance with environmental, non-speech sounds, yet also correlates with speech reception thresholds (Zaar et al., 2023). This suggests that the ACT taps into auditory mechanisms relevant to both speech and non-speech processing, making it a promising clinical tool for assessing auditory function more broadly. Future work could extend this approach using a speech-based version of the change detection task to further clarify how ACT performance relates to real-world speech perception.

While we expected that reduced hearing-in-noise ability would reduce a listeners’ change detection, we also considered the possibility that increased sensitivity to small level changes, often observed in cochlear damage, might facilitate detection. In our previous work (Poole et al., 2025a), when spatial cues were removed, listeners maintained stable performance despite increased scene complexity, suggesting a possible reliance on level cues to detect the appearing source. Individuals with hearing loss, who typically exhibit reduced spatial acuity, might similarly depend more on such level information. To test this idea, we included the SISI test, which measures sensitivity to 1 dB level increments in a pure-tone stimulus and is clinically used to differentiate cochlear from retrocochlear hearing loss. However, we found no effect of SISI performance on change detection. This may reflect the limited variability in our sample where all participants met the clinical criterion for cochlear etiology in both ears for at least 4 kHz. Future work comparing individuals with cochlear and retrocochlear hearing loss could provide a more sensitive test of whether heightened level-change sensitivity contributes to spatial change detection.

Beyond the effects of hearing loss, our findings demonstrated that older age predicted lower hit rates and slower reaction times in detecting the appearing source, indicating reduced accuracy in detecting these spatial changes. Interestingly, false alarm rates were primarily predicted by age rather than hearing metrics, suggesting that age-related changes in attention or decision-making may contribute to response biases independently of auditory sensitivity. To date, little work has examined how aging specifically impacts change detection in complex acoustic scenes. Aging is known to affect cognitive, executive, and attentional functions, with consequences that extend across sensory modalities (Anstey and Low, 2004; Cope et al., 2022; Juan and Adlard, 2019). In the auditory domain, reduced processing resources in older adults may limit the capacity to simultaneously track multiple concurrent objects (Peelle and Wingfield, 2016), diminishing the cognitive resources available for tasks such as change detection (Pronk et al., 2019).

However, it remains unclear why aging gives rise to these difficulties. Although declines in auditory sensory memory have been observed with age, these do not appear to directly affect speech-in-noise performance (Bianco and Chait, 2023), suggesting that aging impacts auditory processing in a task-specific rather than uniform manner. While no prior studies have directly compared younger and older adults using the present change detection paradigm, related work with random tone-pip scenes found that older listeners performed worse than younger ones (de Kerangal et al., 2021). Notably, in that study, change detection ability did not correlate with age or standard audiometric measures (hearing thresholds, speech-in-noise perception, spectral resolution), but instead showed strong associations with auditory sustained attention and musical training. In contrast, we did observe effects of age in the present study, although we did not measure all the factors assessed by de Kerangal et al (2021). This difference between studies may reflect unmeasured covariates of age, such as attentional or cognitive factors, or it may indicate that spatialised change detection engages age-sensitive mechanisms distinct from those involved in non-spatial auditory scene analysis.

### 4.3 Listeners have reduced detection of sound sources from behind them

Consistent with our second research question, listeners showed reduced performance in detecting sounds originating from behind, as reflected by slower reaction times and reduced hit rates. One possible contributor to the delayed detection of sounds behind the listener could be due to the morphology of the head and ears shaping incoming acoustic signals and slightly amplifying frontal sounds while attenuating those originating from behind (Carlile and Pralong, 1994). The degree of this filtering varies between individuals and could, in theory, influence the ease with which appearing sound sources are detected. However, our numerical analysis using head-related transfer functions demonstrated that acoustic shadowing does not explain this deficit, as changes in RMS sound levels for back versus front sources were similar before and after the appearing sound source.

Although we observed small frequency-specific RMS differences of approximately 5 dB, the magnitude of these differences is unlikely to account for the observed reaction time effects. Previous work has shown that reaction time is only weakly influenced by sound level. For instance Kohfeld et al. (1981) found that a 20 dB difference between 60 and 80 phons, levels comparable to those used here (around 62 dB), produced a change in reaction time of only about 15 ms. Similarly, (Kemp, 1984) reported that even a 20 dB change in signal-to-noise ratio yielded reaction time differences of roughly 30 ms. In contrast, our data show reaction time differences of around 200 ms despite maximal frequency-specific RMS differences of just 5 dB. While such effects might be slightly amplified in older listeners, given our demographic, the small level differences observed here are unlikely to explain the magnitude of the reaction time effects.

A more plausible explanation for the delayed reaction times observed in our study is a spatial-attention bias favouring sounds originating from the listener’s forward-facing direction, which aligns with the visual gaze in our task. Previous work has demonstrated that reaction times are fastest when auditory attention is directed toward the same location as gaze (Pomper and Chait, 2017). This alignment may result in slower detection, or “change-deafness”, for sounds occurring outside the visually attended region, while enhancing detection for sounds within visually-attended areas (Best et al., 2023; Demany et al., 2017, p. 201; Eramudugolla et al., 2005; Irsik et al., 2016). Supporting this, recent evidence shows that directing auditory attention to the front or back, even without manipulating gaze direction, can reduce reaction times when detecting amplitude modulation changes, with more pronounced benefits for frontal attention (Bodnár et al., 2024). For hearing-impaired listeners, these spatial-attention mechanisms may interact with reduced auditory resolution or degraded peripheral input, potentially magnifying the difficulty of monitoring less-attended regions.

However, the absence of similar reaction time differences between left and right sources compared to back sources suggests that additional mechanisms may contribute, beyond a forward-facing attentional bias. One possibility is a better-ear effect, whereby lateral sounds produce a higher signal-to-noise ratio at the ear closer to the source. Indeed, we observed an overall RMS difference of approximately 1.5 dB favouring the better ear, which could partially offset the attentional disadvantage associated with being outside the visual field. Alternatively, there may be a gradient of auditory attention extending laterally from the frontal region, though we found no clear evidence for this in the reaction time data. These explanations remain speculative, but they highlight the possibility that different cues may facilitate change detection depending on the source location, for example spatial attention for frontal sounds and acoustic-level cues for lateral ones. Future work could test these possibilities more directly by manipulating spatial attention cues and gaze direction in hearing-impaired listeners, or by increasing the spatial resolution of source locations to examine whether detection varies more finely across near-back positions. Combining such approaches with measures of cognitive load or working memory could further clarify the relative contributions of sensory and attentional factors to spatial change detection.

### 4.4 Conclusion

Together, these findings provide an initial description of how hearing loss, age, and spatial factors interact to shape change detection, and highlight the ACT test as a promising tool for assessing functional hearing beyond audiometric thresholds. However, while we observe significant associations between these measures and performance, the precise mechanisms driving these relationships remain unclear. Future work using physiological measures and targeted manipulations will be needed to disentangle these mechanisms and clarify the causal links between auditory sensitivity, spatial attention, and change detection.

The results nonetheless have implications for hearing aid design and prescription. Noise reduction algorithms can substantially improve speech perception in noise (Jürgens et al., 2025), but potentially at the expense of situational awareness by attenuating non-speech environmental sounds. Our findings suggest that ACT performance may help identify which listeners could tolerate aggressive noise reduction prioritising speech intelligibility versus those who would benefit from preserving background acoustic detail. However, the present study tested listeners without hearing aids, and a critical next step will be to examine how noise reduction processing affects the same individuals, allowing us to quantify the costs imposed by different device settings.

Recent work by Johnson and Healy (2024a, 2024b) demonstrates that balancing speech intelligibility with environmental awareness is achievable by reducing but not eliminating background sound, with benefits for both normal-hearing and hearing-impaired listeners. Incorporating measures like ACT into clinical assessments could help tailor processing strategies to each listener’s perceptual profile. Further research on how noise-reduction algorithms affect spatialised change detection will be essential for optimising hearing-aid fittings that preserve both speech clarity and access to dynamic environmental cues.

## ACKNOWLEDGEMENTS

This study was funded by the William Demant Foundation (Case no. 21-2519).

## 5. AUTHOR CONTRIBUTIONS

**Katarina C. Poole:** Methodology, Software, Formal analysis, Writing - Original Draft, Writing - Review & Editing, Visualization**. Simon With:** Methodology, Investigation**. Vincent Martin:** Methodology, Software**. Maria Chait:** Conceptualization, Writing - Review & Editing, Supervision, Funding acquisition. **Lorenzo Picinali:** Conceptualization, Writing - Review & Editing, Supervision, Funding acquisition**. Martha Shiell:** Conceptualization, Methodology, Formal analysis, Investigation, Writing - Original Draft, Writing - Review & Editing, Supervision, Funding acquisition

## 6. SUPPLEMENTARY

**Table 4:** Table of extended high frequency thresholds for each participant.

| Participant | Left ear |  |  |  | Right ear |  |  |  |
| --- | --- | --- | --- | --- | --- | --- | --- | --- |
|  | 8 kHz | 10 kHz | 12.5 kHz | 16 kHz | 8 kHz | 10 kHz | 12.5 kHz | 16 kHz |
| 1 | 75 | 80 | 85 | NA | 75 | 85 | NA | NA |
| 2 | 75 | 95 | NA | NA | 75 | 95 | NA | NA |
| 3 | 85 | NA | NA | NA | 70 | 85 | NA | NA |
| 4 | 60 | 80 | 75 | NA | 65 | 75 | 80 | NA |
| 5 | 70 | 80 | 85 | NA | 65 | 80 | 85 | NA |
| 6 | 65 | 75 | NA | NA | 70 | 75 | NA | NA |
| 7 | 75 | 85 | NA | NA | 80 | 100 | NA | NA |
| 8 | 60 | 70 | 70 | NA | 65 | 70 | 70 | NA |
| 9 | 75 | 75 | NA | NA | 70 | 75 | NA | NA |
| 10 | 80 | NA | NA | NA | 75 | 80 | NA | NA |
| 11 | 80 | 80 | 90 | NA | 80 | 95 | NA | NA |
| 12 | 65 | 60 | 90 | NA | 60 | 65 | 70 | NA |
| 13 | 60 | 60 | 70 | 60 | 60 | 65 | 80 | NA |
| 14 | 70 | 65 | 75 | NA | 65 | 70 | 85 | NA |
| 15 | 80 | 90 | NA | NA | 65 | 85 | 90 | 60 |
| 16 | 70 | 70 | 75 | 60 | 60 | 60 | 70 | NA |
| 17 | 75 | 85 | 85 | NA | 75 | 75 | NA | NA |
| 18 | 60 | 55 | 60 | 65 | 45 | 45 | 70 | NA |
| 19 | 65 | 80 | 85 | NA | 60 | 70 | 85 | NA |
| 20 | 55 | 50 | 70 | NA | 45 | 45 | 60 | NA |
| 21 | 75 | 85 | 85 | NA | 75 | 90 | NA | NA |
| 22 | 60 | 65 | 70 | 50 | 70 | 65 | 75 | 60 |
| 23 | 55 | 60 | 65 | NA | 50 | 55 | 65 | NA |
| 24 | 55 | 75 | 85 | NA | 50 | 60 | 75 | NA |
| 25 | 60 | 80 | 85 | NA | 70 | 80 | 90 | NA |
| 26 | 75 | NA | NA | NA | 65 | 75 | NA | NA |
| 27 | 55 | 65 | 75 | NA | 65 | 65 | 75 | NA |
| 28 | 45 | 60 | 85 | NA | 40 | 45 | 65 | NA |
| 29 | 65 | 70 | 85 | NA | 70 | 85 | NA | NA |
| 30 | 55 | 65 | 65 | 60 | 55 | 60 | 65 | NA |

**Table 5:** Table of pairwise comparisons.

| **Ear** | **Comparison** | **Frequency Range (Hz)** | **Mean Difference (dB)** | **p-value** | **Max Difference Freq (Hz)** | **Max Difference (dB)** |
| --- | --- | --- | --- | --- | --- | --- |
| **Left** | Front vs Back | 883 - 20000 | 1.585 | 0.0059 | 12808 | 5.54 |
| **Left** | Front vs Left | 20 - 185 | 1.283 | 0.0059 | 20 | 1.651 |
| **Left** | Front vs Left | 289 - 1379 | -1.082 | 0.0059 | 883 | -2.028 |
| **Left** | Front vs Left | 2691 - 10249 | -3.131 | 0.0059 | 6563 | -9.254 |
| **Left** | Front vs Left | 16005 - 20000 | -0.874 | 0.0059 | 20000 | -1.273 |
| **Left** | Front vs Right | 289 - 706 | 0.276 | 0.0117 | 565 | 0.391 |
| **Left** | Front vs Right | 1103 - 20000 | 3.293 | 0.0059 | 12808 | 7.339 |
| **Left** | Back vs Left | 20 - 20000 | -1.349 | 0.0059 | 6563 | -11.752 |
| **Left** | Back vs Right | 706 - 16005 | 1.492 | 0.0059 | 10249 | 4.095 |
| **Left** | Left vs Right | 20 - 20000 | 1.816 | 0.0059 | 6563 | 15.207 |
| **Right** | Front vs Back | 883 - 20000 | 1.606 | 0.0059 | 12808 | 5.394 |
| **Right** | Front vs Left | 20 - 185 | -1.142 | 0.0059 | 20 | -1.356 |
| **Right** | Front vs Left | 289 - 20000 | 2.268 | 0.0059 | 12808 | 7.049 |
| **Right** | Front vs Right | 20 - 148 | 0.869 | 0.0176 | 20 | 1.126 |
| **Right** | Front vs Right | 232 - 1379 | -1.095 | 0.0059 | 883 | -2.295 |
| **Right** | Front vs Right | 4203 - 10249 | -4.861 | 0.0059 | 6563 | -10.185 |
| **Right** | Front vs Right | 16005 - 20000 | -1.122 | 0.0059 | 20000 | -1.346 |
| **Right** | Back vs Left | 20 - 185 | -1.102 | 0.0059 | 20 | -1.388 |
| **Right** | Back vs Left | 706 - 20000 | 1.262 | 0.0059 | 10249 | 3.723 |
| **Right** | Back vs Right | 20 - 20000 | -1.659 | 0.0059 | 6563 | -12.647 |
| **Right** | Left vs Right | 20 - 20000 | -1.913 | 0.0059 | 6563 | -15.847 |
| **Both** | Front vs Back | 452 - 565 | 0.271 | 0.0234 | 565 | 0.343 |
| **Both** | Front vs Back | 883 - 20000 | 1.595 | 0.0059 | 12808 | 5.467 |
| **Both** | Front vs Left | 452 - 4203 | 0.25 | 0.0059 | 3363 | 2.47 |
| **Both** | Front vs Left | 6563 - 16005 | 0.842 | 0.0059 | 12808 | 3.8 |
| **Both** | Front vs Right | 452 - 4203 | 0.22 | 0.0059 | 3363 | 2.462 |
| **Both** | Front vs Right | 6563 - 20000 | 0.474 | 0.0059 | 12808 | 3.767 |
| **Both** | Back vs Left | 452 - 1103 | -0.38 | 0.0059 | 565 | -0.854 |
| **Both** | Back vs Left | 3363 - 20000 | -1.764 | 0.0059 | 6563 | -4.276 |
| **Both** | Back vs Right | 452 - 1103 | -0.468 | 0.0059 | 565 | -0.988 |
| **Both** | Back vs Right | 2691 - 20000 | -1.702 | 0.0059 | 6563 | -4.596 |

